# Deep Genomic Signature for early metastasis prediction in prostate cancer

**DOI:** 10.1101/276055

**Authors:** Hossein Sharifi-Noghabi, Yang Liu, Nicholas Erho, Raunak Shrestha, Mohammed Alshalalfa, Elai Davicioni, Colin C. Collins, Martin Ester

## Abstract

For prostate cancer patients, timing and intensity of therapy are adjusted based on their prognosis. Clinical and pathological factors, and recently, gene expression-based signatures have been shown to predict metastatic prostate cancer. Previous studies used labelled datasets, i.e. those with information on the metastasis outcome, to discover gene signatures to predict metastasis. Due to steady progression of prostate cancer, datasets for this cancer have a limited number of labelled samples but more unlabelled samples. In addition to this issue, the high dimensionality of the gene expression data also poses a significant challenge to train a classifier and predict metastasis accurately. In this study, we aim to boost the prediction accuracy by utilizing both labelled and unlabelled datasets together. We propose Deep Genomic Signature (DGS), a method based on Denoising Auto-Encoders (DAEs) and transfer learning. DGS has the following steps: first, we train a DAE on a large unlabelled gene expression dataset to extract the most salient features of its samples. Then, we train another DAE on a small labelled dataset for a similar purpose. Since the labelled dataset is small, we employ a transfer learning approach and use the parameters learned from the first DAE in the second one. This approach enables us to train a large DAE on a small dataset. After training the second DAE, we obtain the list of genes with high weights by applying a standard deviation filter on the transferred and learned weights. Finally, we train an elastic net logistic regression model on the expression of the selected genes to predict metastasis. Because of the elastic net regularization, some of the selected genes have non-zero coefficients in the classifier which we consider as the DGS gene signature for metastasis.

We apply DGS to six labelled and one large unlabelled prostate cancer datasets. Results on five validation datasets indicate that DGS outperforms state-of-the-art gene signatures (obtained from only labelled datasets) in terms of prediction accuracy. Survival analyses demonstrate the potential clinical utility of our gene signature that adds novel prognostic information to the well-established clinical factors and the state-of-the-art gene signatures. Finally, pathway analysis reveals that the DGS gene signature captures the hallmarks of prostate cancer metastasis. These results suggest that our method helps to identify a robust gene signature that may improve patient management.

## 1 Introduction

Prostate cancer is the most prevalent cancer type among men, with which roughly one in six men will be diagnosed in their lifetime [1]. A majority of detectable prostate cancer cases is entirely quiescent and can be successfully managed without intervention. However, a fraction of prostate cancer cases is aggressive and makes prostate cancer the third leading cause of cancer death among men. This wide range of clinical outcomes makes managing patients with prostate cancer challenging and creates a need to better stratify patients into clinically actionable risk groups. Accurately determining patients’ risk of developing metastasis allows patients with aggressive subtypes to be provided with more intense therapy and protects those with relatively indolent subtypes from the serious side-effects of treatment [2].

Identification of aggressive prostate cancer has been historically achieved through the interpretation of clinical and pathological factors, such as Prostate-Specific Antigen (PSA) and gleason score (it is a score between 2 and 10 for which higher scores mean more advanced cases). Recently, it has been demonstrated that gene expression data within tumor and stroma can also be utilized to identify prostate cancer patients with the risk of developing metastasis [3–6]. For example, Cuzick *et al.* [7] developed a cell cycle proliferation score to identify patients with advanced disease, based on the expression of 31 genes in the cell cycle proliferation pathway. Penney *et al.* [4] developed a gene signature by comparing the gene expression profiles of patients with low gleason scores (≤6) and those with high scores (≥8). Erho *et al.* [8] identified a gene signature to predict early metastasis following radical prostatectomy using a random forest model. Another studies [5, 6] found gene signatures derived from the stroma and microenvironment of the tumor to predict metastasis in prostate cancer.

Since a long follow-up time is needed to determine if a patient is going to develop metastasis, and since high-throughput genomic profiling is costly, all published gene signatures for metastasis were obtained using relatively small labelled datasets (sample size ≤ 545). The combination of a large number of features (genes) and a small number of samples (patients) poses a great challenge to train a classifier to predict metastasis and potentially limits the performance. However, the rise of commercially available high-throughput whole-transcriptome clinical testing, such as the Decipher test (clinicaltrials.gov identifier: NCT02609269), is making large amounts of “unlabelled” data (without metastasis outcome) available.

We hypothesize that such a large unlabelled dataset along with a small “labelled” dataset (metastasis outcome is available) can be utilized to find a better gene signature and as a result train a classifier that predicts metastasis more accurately. We believe a classifier based on such a gene signature can capture prognostic information that cannot be captured by classifiers trained only on the gene signatures obtained from labelled datasets.

To combine unlabelled and labelled datasets for obtaining a gene signature and training a classifier, we want to extract salient features from the gene expressions of the samples in these datasets. Because of the capabilities of deep neural networks to learn features and transfer the learned features between different datasets [9, 10], we adopt an approach based on deep neural networks. Particularly, we utilize Auto-Encoder (AE), a type of unsupervised deep neural networks, that can reduce high-dimensional input data, such as the expression values of all genes, to a low-dimensional representation (salient features) of the input samples. Different variants of AEs have been widely applied for a diverse range of biological problems [10–7]. For example, Denoising Auto-Encoder (DAE)—an extension of AEs which tries to learn salient features from noisy samples—was used to learn complex patterns from gene expression profiles in breast cancer [11]. Similarly, Stacked Denoising Auto-Encoder was shown to be more effective than methods like principal component analysis to obtain functional features from breast cancer gene expression profiles [12]. Therefore, using AEs is a promising approach for our problem. Particularly, since gene expression data are generally noisy, employing DAE is a natural choice to deal with this noise and also to avoid overfitting [18]. Another advantage of using AEs is that transfer learning is applicable on these type of deep neural networks. The features extracted by an AE are transferable between different datasets—generally from a source dataset to a target dataset where the source is larger than the target in terms of the number of input samples [19, 20]. In our context, the source dataset is the large unlabelled dataset, and the target dataset is the small labelled one. Features transferred from the source to the target can be fine-tuned if enough samples are available in the target dataset, or can remain frozen and not trained if the target dataset is small [9].

In this paper, we propose the Deep Genomic Signature (DGS) method to predict metastasis in prostate cancer patients. DGS is based on Denoising Auto-Encoders (DAEs) and transfer learning. We utilize a DAE to extract the most salient features of gene expressions from the samples in a large unlabelled dataset, our source dataset. The learned weights and biases of this DAE are transferred to another DAE, which is trained on a smaller but labelled (with metastasis outcome) dataset, our target dataset. During training of the second DAE, the transferred parameters are frozen, and only the additional parameters are trained. DGS selects the list of genes with high weights by employing a standard deviation filter on the transferred and learned weights of the second DAE. Finally, DGS trains an elastic net logistic regression model [21] on the labelled dataset, using the expressions of the selected genes as features, to predict metastasis. Due to the elastic net regularization, only a subset of the selected genes have non-zero coefficients, and these genes form the DGS gene signature for metastasis in prostate cancer.

We applied DGS to six previously published labelled datasets and one large unlabelled dataset obtained from the Decipher test of GenomeDx Inc. We compared the accuracy of the gene signature discovered by DGS against those of the state-of-the-art signatures for prostate cancer metastasis and showed that the DGS gene signature outperforms the existing signatures in terms of Area Under the receiver operating characteristic Curve (AUC). Further, we performed uni- and multi-variable analyses and found that the DGS gene signature adds novel prognostic information to the well-established clinical factors and the state-of-the-art gene signatures for prostate cancer metastasis. Finally, we performed Gene Set Enrichment Analysis (GSEA) and showed that the DGS gene signature is highly relevant to metastasis and prostate cancer biology. These experimental findings confirmed that our proposed DGS method successfully transfers relevant features from the unlabelled dataset to the labelled dataset. To the best of our knowledge, this is the first study of deep transfer learning from a large unlabelled dataset without clinical outcomes (such as metastasis) to a small labelled dataset with complete clinical outcomes in the context of cancer research. Although we evaluated DGS only in prostate cancer, we believe that DGS is also applicable for other slow progressing cancers for which obtaining (long-term) clinical outcomes requires a lot of time and resources.

## 2 Methods

The proposed DGS method has the following steps (See Figure 1):

1. A DAE is trained on a large unlabelled gene expression dataset to extract salient features of samples in that dataset. More specifically, the DAE is trained to reduce the high-dimensional input of all gene expression values to a low-dimensional representation of a sample. This DAE is the source and we denote it as the *Source*. The output of this step is the learned parameters that extract the most salient features from this dataset. We transfer these learned parameters to the target DAE in the next step.
2. In this step, another DAE is trained on a relatively small labelled dataset. This DAE is the target and we denote it as the *Target*. The *Target* has the transferred parameters from the *Source* and also its additional parameters. Since it is not feasible to retrain the transferred parameters due to the small size of the labelled dataset, they are frozen and not trained, but the additional parameters are trained. We adopt the method of transfer learning from [9]. Transferring parameters from the *Source*—trained on the unlabelled data—to the *Target*—trained on the labelled data—is the core of the DGS method.
3. After training the *Target*, it has the transferred parameters and its additional learned parameters, including the weights of the network. The goal of this step is to define a set of genes whose expressions capture the factors of variation in both the labelled and unlabelled datasets. Such genes provide the classifier (the last step) with enough information to predict metastasis accurately. We select genes with high weights in the *Target* by applying a standard deviation filter on all of the transferred and learned weights. This selection approach is similar to [10–13, 16].
4. The genes selected in the previous step are used to train an elastic net logistic regression model [21] on the labelled dataset to predict metastasis. The genes with non-zero coefficients in the trained model are considered as the DGS gene signature to predict metastasis in prostate cancer.

**Fig. 1.**
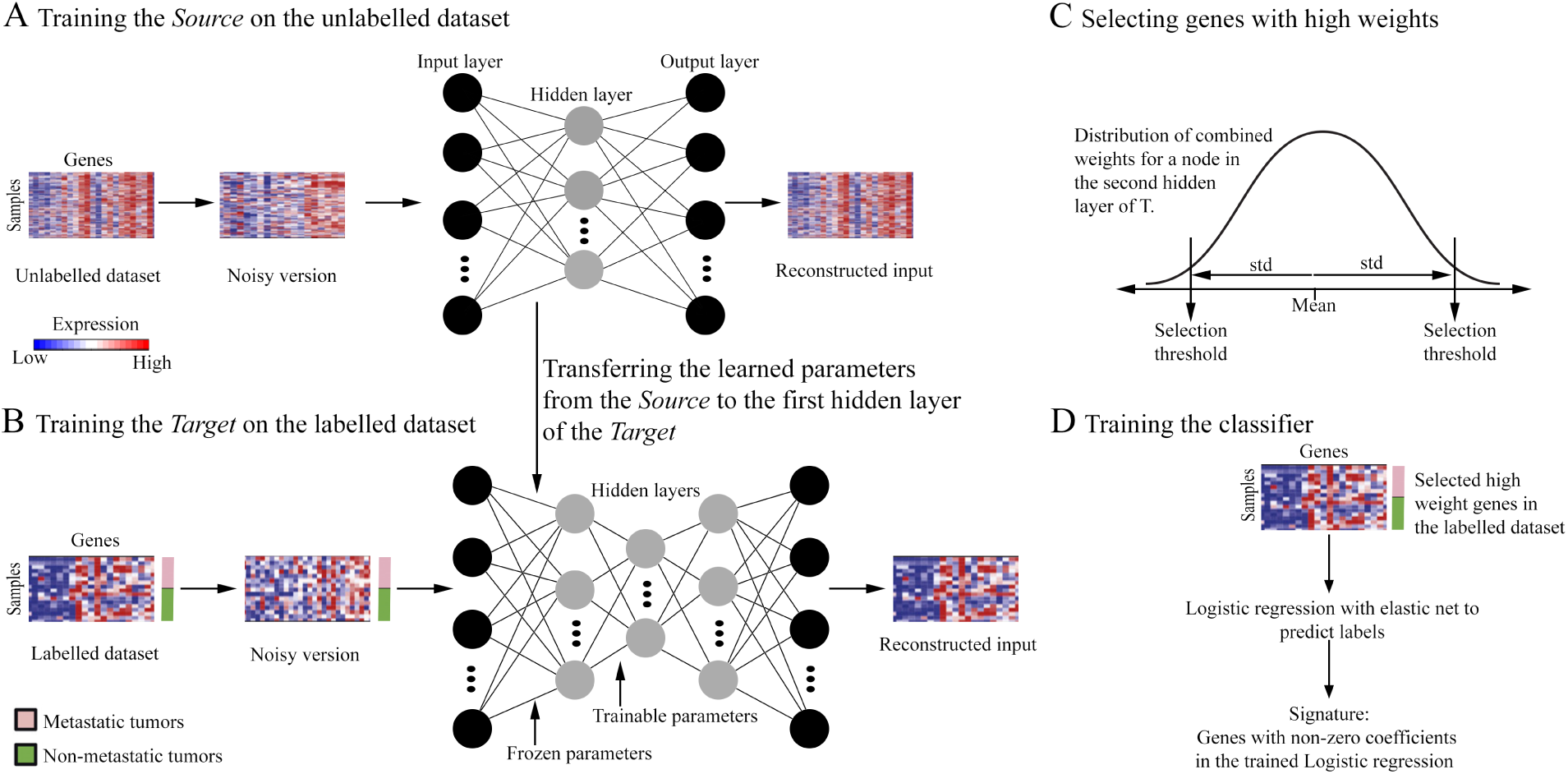
Schematic overview of Deep Genomic Signature. (A) Training the *Source* (a Denoising Auto-Encoder) on the unlabelled data to extract salient features from this dataset. The *Source* has one hidden layer. (B) Training the *Target* (another Denoising Auto-Encoder) on the labelled data. The *Target* has two hidden layers. Parameters of the first hidden layer are transferred from the *Source* which remain frozen and parameters of the second hidden layer (initialized randomly) are trained. (C) Applying a standard deviation filter to select genes based on their weights in the *Target*. These genes are in the tails of the weight distribution of nodes in the second hidden layer of the *Target*. (D) Training an elastic net logistic regression model (*l*_1_ and *l*_2_ regularization) to predict metastasis. The DGS gene signature consists of all of the genes with non-zero coefficients in this classifier.

### 2.1 Extracting features from the unlabelled dataset

The *Source* is a DAE and has three components: an encoder *F* (.), a decoder *G*(.), and a cost function *J* (.). For *M* samples (patients), *N* features (genes) in an unlabelled dataset and *P* nodes in the hidden layer: the encoder takes this unlabelled source dataset *X*_*Source*_ ∈*; ℛ*^*N*^ as input and encodes the expressions of its samples to a lower-dimensional representation *h*_*Source*_∈*; ℛ* ^*P*^. This representation contains the salient features of the input samples. The encoding is performed via a weight matrix *W*_*Source*_ *∈* ∈*; ℛ*^*P*^ and a bias vector *b*_*Source*_ ∈*; ℛ*^*P ×*1^ as follows:

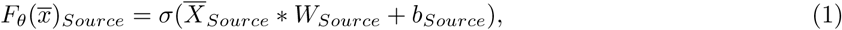

where 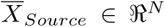 is the corrupted version of input data *X* obtained from a corruption process like 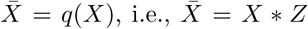, where *Z* is a binary variable sampled from a binomial distribution, and *σ*(.) is the Sigmoid activation function. To use DAE for gene expression data, we need to define a method for corrupting the expression data by adding noise to it. To do so, the values of the expression for some randomly selected patients and genes are changed to zero.

The decoder is a function that maps the learned representation by the encoder back to the input space, attempting to reconstruct the original input:

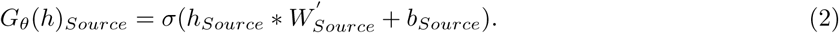

Finally, the following cost function measures the discrepancy between the reconstructed input and the original input (without the added noise):

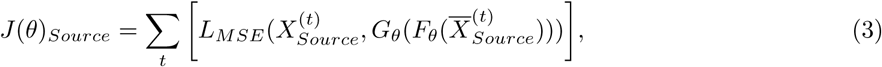

where, *X*^(*t*)^ denotes the input data for sample t and ℒ_*MSE*_(.) is the Mean Squared Error loss function. (.)is the transpose operator because we used the tied weights approach. The set of parameters of the *Source* to be learned and optimized is *θ*_*Source*_ = {*W*_*Source*_, *b*_*Source*_}.

### 2.2 Extracting features from the labelled dataset

The *Target* is another DAE with similar components as the *Source*, but it has two differences compared to the *Source*. The first difference is the network architecture which has two hidden layers instead of one hidden layer in the *Source*. The second difference is that parameters of the first layer, i.e. weights and biases, are not initialized randomly, but these parameters are transferred from the *Source*. These transferred parameters are frozen and not trained during training of the *Target* on the labelled dataset *X*_*Target*_ ∈ *ℛ*^*N*^. The rational behind keeping them frozen is that the number of samples in the labelled data is not sufficient to train both of the layers. Therefore, the parameters of the first layer are not trained, and only the parameters of the second layer are trained:

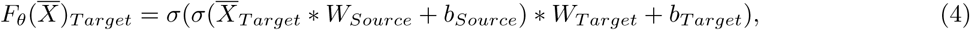

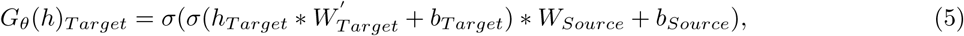

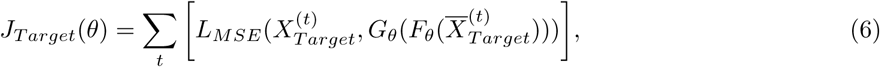

where the notations of symbols and functions are the same as in section 2.1. The set of parameters of the *Target* is *θ*_*Target*_ = {*W*_*Target*_, *b*_*Target*_, *W*_*Source*_, *b*_*Source*_}.

All of the parameters of the *Source* and the *Target* are initialized by the Xavier method [22]. All codes for the *Source* and the *Target* are implemented using the TensorFlow framework in Python 2.

### 2.3 Selecting genes and training the classifier

After the previous steps, we have learned two weight matrices. The first weight matrix is obtained from training the *Source* on the unlabelled data, and the second one is obtained by training the *Target* (only the second hidden layer) on the labelled data. We denote these two matrices as *W*_*Source*_ ∈^*N ×P*^ and *W*_*Target*_ ∈^*P ×Q*^, respectively, where *Q* is the number of nodes in the second hidden layer of the *Target*. We define *W*_*Final*_ ∈^*N ×P*^, as the product of these two matrices as follows:

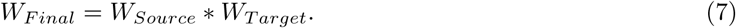

In *W*_*Final*_, each gene is associated with a vector of size *Q* that shows how strongly it contributes to the activation of the nodes in the second hidden layer of the *Target*. The goal of this step is to find a subset of genes that capture the main factors of variation in the gene expression of the samples in labelled and unlabelled datasets, i.e. the genes with high weights in *W*_*Final*_. To select these genes, we apply a standard deviation filter on the weights of the *Q* nodes in *W*_*Final*_. This filter determines the genes whose values fall into the tails of the weight distribution—regardless of the sign—for each node. Since this method is unsupervised, some of these genes may be irrelevant for predicting metastasis. After selecting the genes with high weights, their expression is used as features for an elastic net logistic regression [21] model to predict metastasis. After training this classifier on the labelled dataset, we select those genes (features) with non-zero coefficients in the classifier as the DGS signature. We note that these two phases of gene selection are essential because the genes selected by the standard deviation filter are not necessarily associated with metastasis. However, all of the genes with non-zero coefficients in the classifier are associated with metastasis.

Training and hyper-parameter tuning of the gene selection and classifier are implemented in R 3.4.

### 2.4 Datasets

A total of 17,352 de-identified and anonymized radical prostatectomy prostate cancer tumor gene expression profiles using the Human Exon 1.0 ST microarray (ThermoFisher, Carlsbad, CA) are retrieved from the Decipher GRID. This include gene expression data from six published retrospective cohorts (sample size = 2, 216) with complete long-term outcomes along with their metastasis status (we refer to these cohorts as the labelled datasets) and 1 prospective cohort (sample size = 15, 136) with basic pathological information but without metastasis status (we refer to this cohort as the unlabelled dataset). Labelled datasets are as follows: Mayo I and II (GSE46691 and GSE62116) [8, 23, 24]; CCF (GSE62667) [25, 26]; JHMI (GSE79957 and GSE79956) [27]; TJU (GSE72291) [28]; MetaSet [29]^1^. The unlabelled dataset is obtained from clinical use of the Decipher test (GenomeDx Inc., San Diego, CA; clinicaltrials.gov identifier: NCT02609269). The microarrays are normalized using the Single Channel Array Normalization (SCAN) algorithm and summarized by gene annotation [30]. The normalized expression data have 46,050 features (genome-wide) and all of the datasets use the same number of features. The number of features is reduced to 13,891 by excluding features with low levels of detection and signal. Batch effect is considered and carefully monitored for all of the utilized datasets by checking the heatmap/hierarchical clustering to make sure that samples do not cluster together based on their cohorts.

### 2.5 Clinical evaluation

We evaluate performance of DGS against five state-of-the-art baselines: Erho *et al.* [8], Penney *et al.* [4], Cuzick *et al.* [7], Mo *et al.* [5], and Tyekucheva *et al.* [6] in terms of AUC to predict metastasis on five validation datasets. We use the Delong method [31, 32] to calculate confidence interval. We further assess the clinical utility of the DGS gene signature with survival analyses on the MetaSet. We use uni-variable and multi-variable Cox proportional hazard models with mixed effects (CoxPHME) [33] to assess the associations of the gene signature obtained by the DGS method with the time to metastasis. In order to demonstrate the independent prognostic power of the obtained gene signature, we use the multi-variable analysis and adjust for the common clinical and pathological factors that have been shown to be prognostic in prostate cancer. In addition, we run multi-variable CoxPHME with the gene signature obtained by DGS adjusting first for each baseline separately and then for all of the baselines combined. We divide the prediction scores from all of the five baselines^2^ as well as the DGS by 0.1, such that the reported Hazard Ratio (HR) should be interpreted for each 10% increase in the predictor score.

### 2.6 Biological evaluation

We investigate the pathway and functional enrichment for the genes that have non-zero coefficients in the classifier. We perform a hypergeometric test based Gene Set Enrichment Analysis (GSEA) (*https://github.com/raunakms/GSEAFisher*). Gene sets of pathways are obtained from Molecular Signature Database v6.0 [34]. A cut-off threshold of false discovery rate < 0.01 is used to obtain the significantly enriched pathways. We select only gene sets/pathways with at least three genes enriched from the query list. Analysis for biological and clinical evaluations of this gene signature is performed in RStudio 1.0.143. All codes are publicly available: https://github.com/hosseinshn/Deep-Genomic-Signature

## 3 Results

### 3.1 Experiments

In our experiments, we investigate the following questions:

1. Does DGS that uses both unlabelled and labelled datasets outperform the baselines that only use labelled datasets in terms of AUC to predict metastasis?
2. Does DGS capture novel prognostic information that well-established clinical factors such as PSA and gleason score cannot capture?
3. Does DGS capture novel prognostic information that the baselines cannot capture?
4. Does the gene signature discovered by the DGS associate with metastasis and prostate cancer biology? To answer these questions, we train the DGS in the following steps: training the Source on the unlabelled data, training the *Target* on the Mayo I dataset—the largest labelled dataset—using the learned parameters transferred from the *Source*, selecting genes with high weights in the *Target* as described in the method section, and training the classifier on Mayo I to predict metastasis. To answer the first question, we train all of the baselines on the same labelled dataset and compare their performance (AUC) to DGS. To answer the second and third questions, we employ uni-variable and multi-variable analyses to study associations of the DGS method, the clinical factors, and the baselines with time to metastasis. Finally, to answer the fourth question, we apply GSEA for the genes with non-zero coefficients in the classifier part of DGS and study the relevance of the enriched pathways to metastasis.

#### Training the *Source*

In order to train the *Source*, three randomly selected sample subsets of size 12000, 2570, and 566 were dedicated to train, development, and test sets, respectively. The *Source* was trained using the train set, validated on the development set, and tested with the unseen test set. For the *Source*, we investigated diverse sets of hyper-parameters to determine the size of hidden layer, learning rate, batch size, corruption rate, and number of epochs. We minimized the cost function by Adagrad optimization method [35] and selected the following values for the hyper-parameters in the *Source*: 256 nodes in the hidden layer (from a set of 256, 512, 1024, 2048, and 4096); learning rate of 0.05 (from a set of 0.05, 0.01, and 0.1); batch size of 100 (from a set of 10, 100, 500, and 1000); corruption level of 0.2 (from a set of 0.1, 0.2, 0.3 and 0.5); and number of epoch was set to be 1000 (from a set of 100, 200, 300, 500, and 1000). Obtained costs for train, development, and test for the *Source* were 0.007, 0.008, and 0.008, respectively.

#### Training the *Target*

We used Mayo I dataset to train the *Target*. A set of 359 randomly selected samples were dedicated to train, and the rest for development and test (186 samples) similar to [8]. We selected the following values for hyper-parameters of the *Target* after tuning: 64 nodes in the second hidden layer of the *Target* (selected from a set of 32, 64, and 128); learning rate of 0.01 (from a set of 0.01, 0.05, and 0.1); batch size of 359 (from a set of 5, 10, 50, 100, and 359); number of epochs of 1000 (from a set of 300, 500, and 1000). Although we examined the same corruption rates as for the *Source*, the input data without noise had the best performance. Activation function, cost function, and optimizer were also the same as the *Source*. Train and test costs for the *Target* were 0.031 and 0.036, respectively.

#### Selecting genes

In order to select genes with high weights, we set the threshold to be 3.94 which means only genes whose weights are 3.94 standard deviations smaller or greater than the mean (average value of all of the input weights of a node) are selected to be the input of the classifier. In the previous studies the value of this threshold for similar purposes was typically set to 2 [11–13] or 2.5 [10, 16]. Possibly due to different ranges of the weights or different normalization, these typical thresholds resulted in excessive number of features (genes) on a rather small training dataset. We applied the following backward selection approach: we started from 4 and decreased the value of the threshold by steps of 0.01, observed a jump in the number of selected genes at the value of 3.94, and chose it as the threshold. This threshold was chosen without considering prior knowledge about the biology of the selected genes and without training separate classifiers for the considered thresholds. By applying this threshold, we obtained 141 genes. This value forthe threshold provided us with a fair comparison to the state-of-art signatures as their number of features range from 22 to 157. We further used these genes to train an elastic net logistic regression model to predict metastasis.

#### Training the classifier

To study the performance of the selected genes to predict metastasis, an elastic net logistic regression model (*l*_1_ and *l*_2_ regularization) was trained on Mayo I (with same train and validation split). Results (training and validation AUCs) for training the classifier are available in Table S1. Two hyper-parameters *γ* > 0, the complexity parameter, and 0 *≤ α ≤*1, the compromise between *l*_2_ (*α* = 0) and *l*_1_(*α* = 1) penalties, were optimized under cross validation (*α* = 0.6 and *γ* = 0.032). 38 genes had non-zero coefficients in the logistic regression model (Table S2). Scores from this model predict the probability of a prostate cancer patient developing metastasis.

### 3.2 Clinical evaluation

#### DGS predicts prostate cancer metastasis

In the blinded validation we found that the DGS method is predictive of the metastatic disease across five external validation datasets (CCF, Mayo II, JHMI, TJU, and MetaSet) with AUCs ranging from 0.67 to 0.83 (Table 1). Our gene signature outperformed all previously published gene signatures in four validation datasets (CCF, Mayo II, TJU, and MetaSet). Erho *et al.*’s gene signature was found to be the most predictive classifier in JHMI. Likewise, Mo *et al.*’s gene signature tied DGS for highest AUC in Mayo II. Erho *et al.*’s gene signature had the second highest AUC in three datasets (Mayo II, TJU, and MetaSet). Penney *et al.*’s gene signature was the second most predictive classifier in JHMI and MetaSet (tied with Erho *et al.*). Mo *et al.*’s gene signature had the second best performance in one testing dataset (CCF). Penney *et al.*’s gene signature had the third highest AUC in three datasets (CCF, Mayo II, and TJU). Cuzick *et al.*’s gene signature had the third highest AUC in MetaSet cohort. Likewise, the DGS gene signature was the third most predictive classifier in JHMI testing dataset. Among the 38 genes with non-zero coefficient in the classifier, we only had six genes in common with the compared state-of-the-art gene signatures (one with Erho *et al.* and five with Penney *et al.*). This indicates that the added genes by the DGS provided contributions to the model and improvement in the performance over the other gene signatures.

**Table 1.** Comparison of signatures based on AUC (with confidence interval) of prediction of metastasis on five independent validation datasets.

| Study/Dataset | CCF | Mayo II | JHMI | TJU | MetaSet |
| --- | --- | --- | --- | --- | --- |
| Erho *et al.* [8] | 0.78 (0.70-0.85) | 0.74 (0.67-0.81) | **0.73 (0.67-0.79)** | 0.74 (0.61-0.87) | 0.70 (0.64-0.76) |
| Penney *et al.* [4] | *0.80 (0.74-0.87)* | *0.71 (0.65-0.78)* | 0.72 (0.65-0.78) | *0.68 (0.53-0.83)* | 0.70 (0.65-0.76) |
| Cuzick *et al.* [7] | 0.73 (0.65-0.81) | 0.64 (0.57-0.72) | 0.63 (0.55-0.70) | 0.67 (0.51-0.82) | *0.67 (0.60-0.73)* |
| Mo *et al.* [5] | 0.81 (0.74-0.88) | **0.79 (0.73-0.85)** | 0.58 (0.51-0.66) | 0.59 (0.35-0.83) | 0.65 (0.59-0.72) |
| Tyekucheva *et al.* [6] | 0.71 (0.62-0.79) | 0.64 (0.56-0.71) | 0.63 (0.56-0.70) | 0.55 (0.39-0.72) | 0.58 (0.51-0.64) |
| DGS | **0.83 (0.76-0.89)** | **0.79 (0.73-0.85)** | *0.67 (0.61-0.74)* | **0.79 (0.69-0.88)** | **0.73 (0.68-0.79)** |

All methods are trained on Mayo I; **boldface** indicates the best method of that corresponding dataset; underline indicates the second best method(s) for that dataset, the third best method(s) are indicated in *italic* form.

#### DGS captures novel prognostic information in addition to the clinical factors

In order to evaluate DGS for clinical utility, we preformed uni-variable and multi-variable survival analyses in the Metaset against different clinical risk factors (Table 2). uni-variable revealed that, except for surgical margins and PSA, DGS (HR 1.23, p < 0.001) along with other factors were significantly associated with metastasis. Likewise, in multivariable analysis, when DGS was adjusted for the same clinical factors, DGS still remained significantly associated with metastasis (HR of 1.14, *p*-value<0.001). This observation indicates that DGS captured additional prognostic information beyond these standard clinical and pathological factors.

**Table 2.**
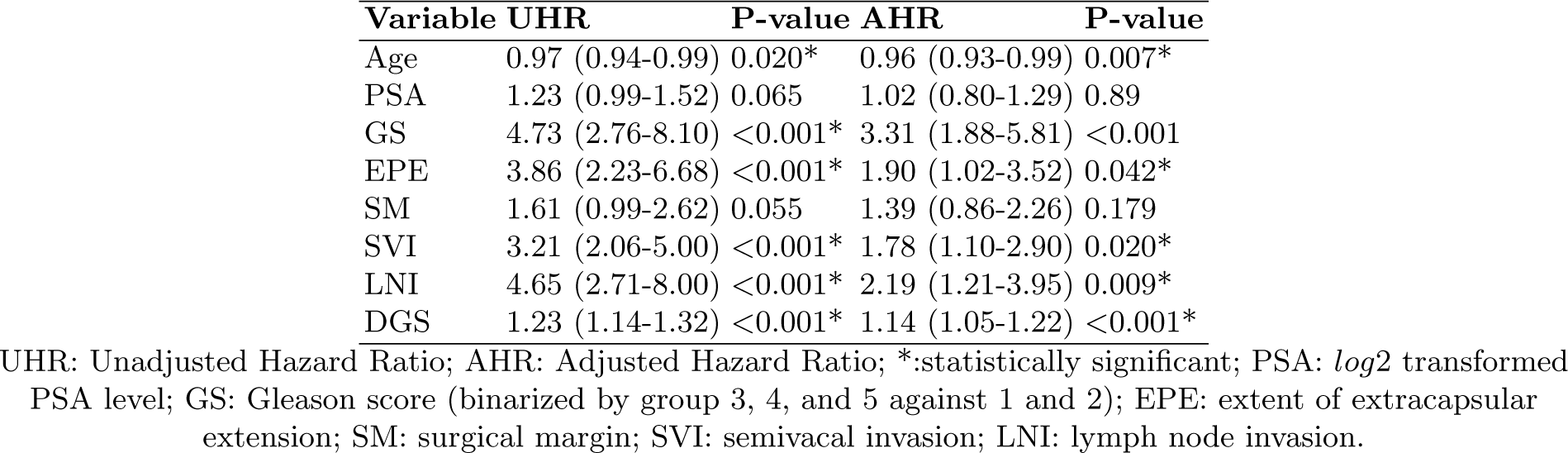
Uni-variable and multi-variable Cox Model with random effects on the MetaSet.

#### DGS captures novel prognostic information in addition to the baselines

We also performed multivariable survival analyses of the DGS with each of the previously published gene signatures individually. When assessed against these five signatures separately, DGS was found to be significant in all of them and which means it contained independent prognostic information not captured by the previously published gene signatures (Table 3). Furthermore, in order to determine if DGS captured novel prognostic information beyond what was captured by the previously published gene signatures combined, we performed a single multi-variable analysis of DGS adjusting for all of the published genomic signatures (6 variables in total). We found that again DGS remained significant indicating it captured prognostic information that no other gene signature had captured (Table 4).

**Table 3.** Multi-variable analysis adjusting other baselines as well as clinical variables on Meta Set.

| Method | AHR | P-value |
| --- | --- | --- |
| Erho *et al.* [8] | 1.21 (1.06-1.38) | 0.006* |
| DGS | 1.09 (1.00-1.18) | 0.039* |
| Penney *et al.* [4] | 1.42 (1.22-1.65) | <0.001* |
| DGS | 1.14 (1.06-1.23) | <0.001* |
| Cuzick *et al.* [7] | 1.09 (0.95-1.25) | 0.235 |
| DGS | 1.10 (1.01-1.20) | 0.024* |
| Mo *et al.* [5] | 1.04 (0.98-1.11) | 0.178 |
| DGS | 1.12 (1.04-1.21) | 0.004* |
| Tyekucheva *et al.* [6] | 0.99 (0.89-1.10) | 0.825 |
| DGS | 1.14 (1.05-1.23) | <0.001* |

AHR: Adjusted Hazard Ratio;(.):Confidence Interval;*:statistically significant

**Table 4.** DGS multi-variable analysis adjusting for five published gene signatures.

| Methods | AHR | P-value |
| --- | --- | --- |
| Erho *et al.* [8] | 1.16 (1.01-1.33) | 0.040* |
| Penney *et al.* [4] | 1.16 (1.00-1.34) | 0.048* |
| Cuzick *et al.* [7] | 1.35 (1.16-1.57) | <0.001* |
| Mo *et al.* [5] | 1.04 (0.97-1.10) | 0.261 |
| Tyekucheva *et al.* [6] | 1.00 (0.91-1.11) | 0.951 |
| DGS | 1.10 (1.01-1.20) | 0.031* |

AHR: Adjusted Hazard Ratio;(.):Confidence Interval;*:statistically significant

### 3.3 Biological evaluation

#### Enriched pathways for the DGS gene signature

We used GSEA to find the signaling pathways regulated by the 38-gene signature with non-zero coefficients (Figure 2-A). We obtained pathways such as Muscle Contraction (*p* = 7×10^-8^) and Actin-Cytoskeleton Regulation (*p* = 6×10^-5^) as the top-hits. These pathways are critical for the migration of metastatic initiating cells from the primary tumor to the circulatory system invading the stroma [36, 37]. The enriched pathways corroborate the importance of stroma in metastasis [38]. This 38-gene signature was also enriched for Androgen Response pathway (*p* = 4×10^-5^) that is critical for prostate cancer progression [39].

**Fig. 2.**
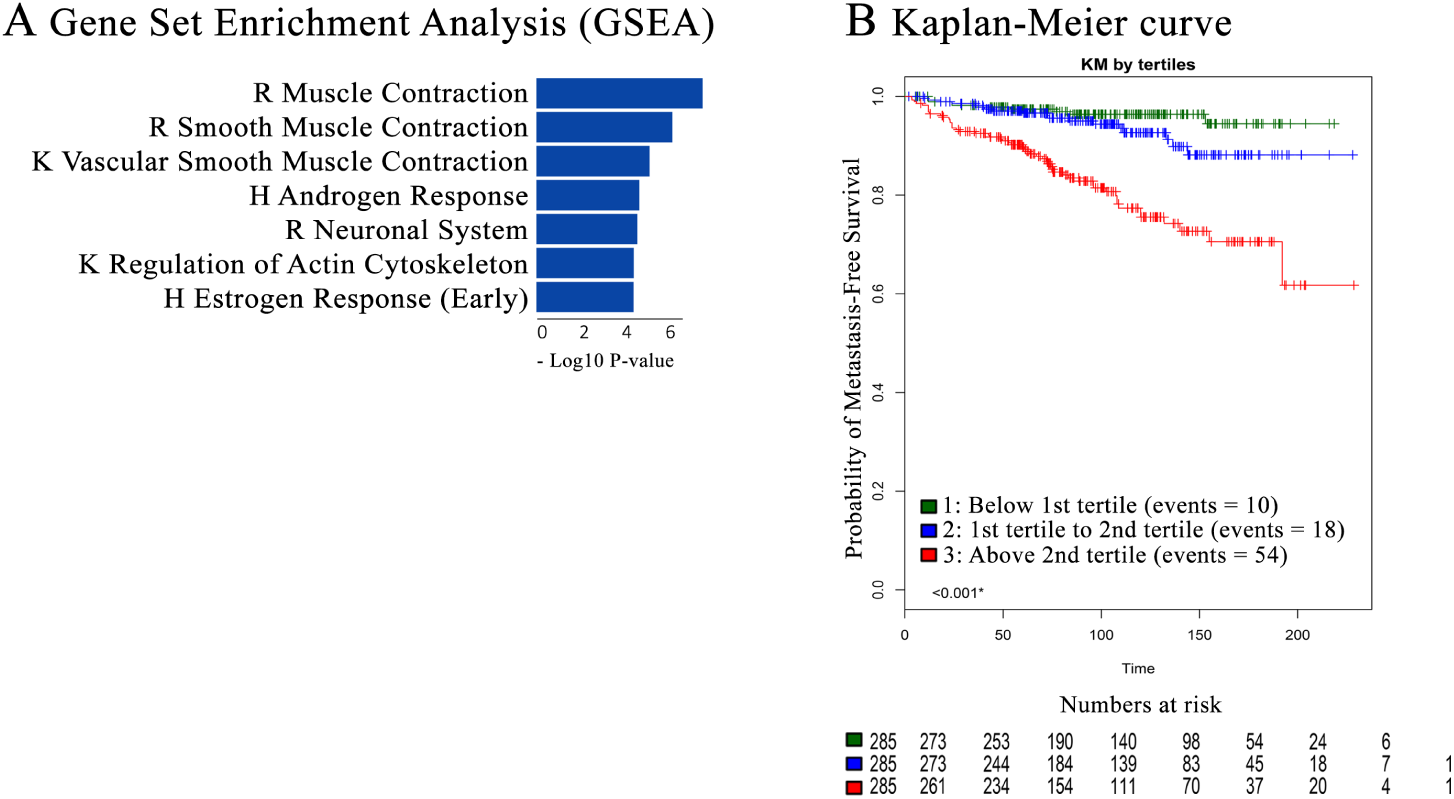
Biological evaluation of the gene signature obtained by DGS method. (A) GSEA for the genes with non-zero coefficients in the classifier (K, R, and H denotes KEGG, REACTOME, and Hallmarks resources, respectively). (B) KM curve for the DGS gene signature that indicates it can successfully distinguish indolent cases from metastatic ones.

#### Kaplan-Meier survival

We also performed a Kaplan-Meier (KM) survival analysis for the DGS results. The KM curve by tertiles in MetaSet [29] demonstrated that DGS can successfully distinguish primary prostate cancer tumors with metastatic potential from indolent tumors (Figure 2-B).

## 4 Discussion

In this paper, we investigated the problem of gene signature discovery for prostate cancer metastasis. We addressed the challenge of small labelled train datasets (with metastasis outcome) by utilizing an unlabelled dataset (without metastasis outcome) together with the labelled ones. We proposed to use DAE and transfer learning to extract features from the expressions of samples in a large unlabelled dataset and a small labelled one. These extracted features were used in the form of genes with high weights to train an accurate elastic net logistic regression model to predict metastasis. We found a novel gene signature based on those genes that had non-zero coefficients in this classifier. To our knowledge, Deep Genomic Signature (DGS) is the first method that learns features from unlabelled and labelled datasets together to predict an outcome (metastasis) in cancer research. Although in this paper we only focused on prostate cancer and metastasis, in practice, the proposed method can work with other cancers and outcomes (phenotype) as well.

The prognostic capability of the discovered gene signature by DGS method was successfully validated on five independent datasets and showed superior performance compared to the state-of-the-art genomic signatures for prostate cancer metastasis. More importantly, the gene signature obtained by the DGS provided independent prognostic information in addition to the clinical and pathological factors. Moreover, DGS gene signature also provided novel information to each and all of the state-of-the-art genomic signatures (separately and combined) according to multi-variable analyses. This observation was indicative of effectiveness of the proposed gene signature for prognostic purposes. Genes that had non-zero coefficients (in the classifier) in the gene signature obtained by the DGS method were enriched for signaling pathways highly relevant to prostate cancer metastasis. The gene signature included novel genes such as PGM5P4-AS1 that have not been associated with prostate cancer metastasis before and thus require further experimental validation.

In addition to the proposed method, we experimented with several other alternative methods based on DAEs. For example, we attempted to train a classifier on the learned representation of the *Target* (*h*_*Target*_). To our surprise, this model was not able to predict metastasis accurately (AUC=0.64 on the Mayo I test set instead of AUC=0.72 of the current model). We also considered nonlinear classifiers such as multilayer neural network on this representation, but did not observe a significant improvement. From the computational point of view, one possible explanation is that the learned representation had not converged to a stable state. We argue that if this was the case, we would not have been able to train an accurate logistic regression model based on high weight genes that are essentially extreme cases in capturing factors of variation in the train dataset. We also trained another logistic regression model based on high weight genes obtained from only the *Source* (the first DAE trained on the unlabelled dataset) and its performance was still fairly competitive. Another explanation is that training the *Target* distorts the salient features extracted from the unlabelled dataset. We analyzed this by training two logistic models based on learned representations of the *Source* (*h*_*Source*_) and the *Target* (*h*_*Target*_) separately and observed an increment in the AUC (from AUC=0.60 for the *Source* to AUC=0.64 for the *Target*). This observation indicated that the *Target* made the learned representation more suitable to predict metastasis. We believe the explanation for the poor performance of the classifier based on the learned representation is that the learned factors of variation by DAEs were associated with some other unobserved phenotype(s) (other than metastasis). In the abstract representation level, signals that were associated with metastasis were distorted with strong factors of variation associated with that unobserved phenotype(s). On the other hand, this issue was less severe at the input (genes) level because we only focused on those genes that had high weights and were more distinguished than the other genes. These high weight genes captured most of the factors of variations in both of the unlabelled and labelled datasets with less distortion by the other genes. For future direction, this problem can be explored from a deep semi-supervised learning point of view [40].

## 5 Conclusion

In this paper, we introduced the Deep Genomic Signature (DGS) method based on Denoising Auto-Encoders (DAEs) and transfer learning. DGS bridges the gap between an unlabelled dataset and a labelled one to train an accurate prognostic model for prostate cancer metastasis prediction. This method can be adapted to a wide range of cancers that are populated with unlabelled samples. Using DGS, we discovered a gene signature predictive of prostate cancer metastasis. This demonstrates that including unlabelled gene expression data along with the labelled data in signature discovery, improves the accuracy to predict metastasis. Overall, our approach helps to identify a robust gene signature that may improve patient management.

## Acknowledgements

We would like to thank our collaborators from Johns Hopkins (Ashley Ross, Bruce Trock, and Edward M. Schaeffer), Mayo Clinic (Robert B. Jenkins and R. Jeffrey Karnes), Thomas Jefferson University (Robert Den, Adam Dicker, Leonard Gomella), Cleveland Clinic (Eric Klein), and others at these institutions for supporting this work by providing patients data with clinical annotations and insights into their interpretation. We would also like to thank Yen-Yi Lin from the Vancouver Prostate Centre for his support.

## Funding

This research was supported by Canada Foundation for Innovation (33440 to C.C.C.), The Canadian Institutes of Health Research (PJT-153073 to C.C.C.), Terry Fox Foundation (201012TFF to C.C.C.), The NSERC CREATE program, Computational Methods for the Analysis of the Diversity and Dynamics of Genomes (To H.SN. and M.E.), and a Discovery Grant from the National Science and Engineering Research Council of Canada (to H.SN. and M.E.).

## Conflict of interests

Y.L., N.E., M.A., and E.D. were employees of GenomeDx Inc. at the time the study was performed.

## Appendix

**Table S1:** Train AUCs (confidence interval) on Mayo I dataset for all of the methods.

| Method/data | Mayo I-train | Mayo I-validation |
| --- | --- | --- |
| Erho et al. | 0.909 (0.879-0.938) | 0.741 (0.666-0.815) |
| Penney et al. | 0.748 (0.698-0.799) | 0.701 (0.624-0.778) |
| Cuzick et al. | 0.539 (0.478-0.6) | 0.453 (0.369-0.538) |
| Mo et al. | 0.923 (0.896-0.951) | 0.61 (0.526-0.693) |
| Tyekucheva et al. | 0.802 (0.757-0.848) | 0.61 (0.527-0.693) |
| DGS | 0.823 (0.781-0.866) | 0.726 (0.649-0.802) |

**Table S2:** Genes with non-zero coefficients in the DGS classifier.

| Marker |
| --- |
| A2M |
| ABHD2 |
| ANTXR2 |
| AZGP1 |
| C1QTNF9B |
| C1QTNF9B-AS1 |
| CACNA1D |
| CALD1 |
| CANX |
| ENDOD1 |
| ERG |
| FAM13C |
| FAM19A5 |
| GREB1 |
| GSTM2 |
| HS6ST1 |
| INPP4B |
| IRS4 |
| ITGA1 |
| KCNMB1 |
| KCNN4 |
| LCE3B |
| MEIS2 |
| MYBPC1 |
| MYL12B |
| MYO6 |
| NCAM2 |
| NRAP |
| ODAM |
| PGM5P4-AS1 |
| PLS3 |
| RLN1 |
| RLN2 |
| RP11-33A14.1 |
| SCIN |
| SPON2 |
| SYT7 |
| TENM1 |

## Footnotes

1 A subset of patients in this cohort were included in Mayo II, JHMI, TJU cohorts.

2 prediction scores of [4] and [7] are scaled to (0, 1) first.

